# The effect of paracrine factors released by irradiated peripheral blood mononuclear cells on neutrophil extracellular trap formation

**DOI:** 10.1101/2022.05.30.493953

**Authors:** Katharina Klas, Anna S Ondracek, Thomas M Hofbauer, Andreas Mangold, Karin Pfisterer, Maria Laggner, Dragan Copic, Martin Direder, Daniel Bormann, Hendrik Jan Ankersmit, Michael Mildner

## Abstract

Neutrophil extracellular trap (NET)-formation represents an important defence mechanism for rapid clearance of infections. However, exaggerated NET formation has been shown to negatively affect tissue-regeneration after injury. As our previous studies revealed strong tissue-protective and regenerative properties of the secretome of stressed peripheral blood mononuclear cells (PBMCsec), we here investigated the influence of PBMCsec on the formation of NETs. The effect of PBMCsec on NET formation was assessed ex vivo in ionomycin stimulated neutrophils derived from healthy donors using flow cytometry, image stream analysis and quantification of released extracellular DNA. Molecular mechanisms involved in NET formation that were potentially impaired by PBMCsec treatment, including protein kinase C activity, reactive oxygen species production and peptidyl arginine deiminase 4 activity were analysed. Our results showed that PBMCsec significantly inhibited NET formation. Investigation of the different biological substance classes found in PBMCsec revealed only partial reduction of NET formation, suggesting a synergistic effect. Mechanistically, PBMCsec treatment did not interfere with calcium signalling and PKC-activation, but exerted anti-oxidant activity, as evidenced by reduced levels of reactive oxygen species and upregulation of heme oxygenase 1, hypoxia inducible-factor 1 as well as heat shock protein 27 in PBMCsec-treated neutrophils. In addition, PBMCsec strongly inhibited the activation of peptidyl arginine deiminase 4 (PAD4), ultimately leading to the inhibition of NET formation. As therapeutics antagonizing excessive NET formation are currently not available, our study provides a promising novel treatment option for a variety of conditions resulting from exaggerated NET formation.

## Introduction

Neutrophil granulocytes represent the main population of circulating leukocytes in the blood (1). They exert a plethora of functions critical for maintaining immune homeostasis and their contribution to immune regulatory mechanisms is of vital importance during infectious conditions (2). Being amongst the first cell populations to be recruited to a site of infection, they use a broad machinery of defence mechanisms including the production of reactive oxygen species (ROS), excretion of cytotoxic granules, phagocytosis of pathogens and the formation of neutrophil extracellular traps (NETs) to fight invading pathogens (3). Besides these functions, NET formation represents another potent defence mechanism for the elimination of pathogens (4). Neutrophils are equipped with a vast array of surface receptors, including Toll-like and NOD-like receptors, G-protein coupled receptors, cytokine receptors as well as Fc and complement receptors, rendering them highly responsive to a multitude of stimuli (5–9). Some of these stimuli, such as ROS or bacterial toxins, are potent inducers of NETs and operate independently of the classical neutrophil activation pathways via surface receptors (10–12). After induction of NET formation, intracellular Ca^2+^ levels increase due to influx and its release from the endoplasmic reticulum, promoting protein kinase C (PKC) activation and phosphorylation of Gp91^phox^ (13). Activation of PKC, in turn, leads to the assembly of functional NADPH oxidase, which generates reactive oxygen species (ROS) (14). Furthermore, increased Ca^2+^ levels activate peptidyl arginase deiminase 4 (PAD4) (15), which promotes chromatin de-condensation by converting arginine residues of core histones H3 and H4 into citrulline (12). In addition, ROS also lead to the gradual disassembly of the nuclear membrane, followed by the dispersion of chromatin throughout the cytoplasm, where it is decorated with granular and cytoplasmic contents (16). Ultimately, chromatin, DNA, granular and cytoplasmic contents are released into the extracellular space as NETs (10,17).

Neutrophil functions, specifically the extrusion of NETs, are considered beneficial during infection (18). However, dysregulated or extensive NET formation may result in undesirable tissue damage (19,20) and is linked to many inflammatory disorders, including sepsis, asthma, lupus, rheumatologic diseases as well as diabetes (21). Additionally, neutrophils receive increasing interest in cancer research as potential drivers of metastasis (22). Furthermore, NETs are discussed as potential inducers of endothelial tissue damage, leading to various forms of vasculitis (23). The accumulation of neutrophils, as well as the entailing activation and NET formation, at the culprit site lesion during acute coronary syndrome or acute myocardial infarction is associated with poor disease prognosis and an increased long-term mortality rate (24–26)(25,26)(25,26). In addition to systemic disorders, excessive NET formation is also associated with locally impaired or prolonged tissue regeneration, due to increased neutrophil-derived ROS in the microenvironment of the injury (20,27–30).

Recent advances in cell-derived, yet cell-free medicinal products have increasingly gained attention in regenerative medicine (31,32). While initial research on cell-free therapeutic agents focused on secretomes derived from stem cells, we could demonstrate that the secretome of peripheral blood mononuclear cells (PBMC) exhibits comparable regenerative effects (33,34,43–46,35–42). The potency of the PBMC-derived secretome (PBMCsec) was further increased by exposing PBMCs to 60 Gy γ-irradiation, which induces apoptosis and necroptosis, resulting in the release of a plethora of pro-regenerative paracrine factors (33). Lichtenauer et al. showed strong regenerative potential of PBMCsec in rodent and porcine models of acute myocardial infarction (37,47). These pioneering findings laid the foundation for further studies, which identified a broad spectrum of therapeutic implications for PBMCsec in a vast variety of pathologic conditions, including chronic heart failure after myocardial infarction (48), cerebral ischemia (44), burn injury (43), diabetic wound healing (34), and acute spinal cord injury (45). Furthermore, strong anti-inflammatory properties of PBMCsec have been demonstrated in the context of myocarditis (38) as well as inflammatory skin conditions (41). The observed tissue-regenerative effect of PBMCsec is based on a complex interplay of various biologically active agents produced and released by stressed PBMCs (34,35,37). The broad action spectrum of PBMCsec has been intensively investigated and revealed promising treatment opportunities, where anti-inflammatory (49,50), anti-microbial (36), tissue-regenerative (35), pro-angiogenic (34,51) and vasodilatory (40) properties are important.

Although PBMCsec possesses compelling immunomodulatory effects (38), potential anti-inflammatory and stabilizing effects on (activated) neutrophils have not been investigated so far. Hence, we sought to investigate the effect of PBMCsec on NET formation.

## Results

### PBMCsec inhibits NET formation

Although immunomodulatory properties of PBMCsec have been well described in a variety of different cell types (34,41,43), a potential effect of PBMCsec on neutrophils has not yet been explored. To investigate whether PBMCsec interferes with experimentally induced NET formation, we pre-incubated human whole blood with 2 units/mL PBMCsec or an equivalent dose of vehicle prior to stimulation with 5 μM ionomycin. Unstimulated or PBMCsec-treated samples showed few citH3-positive neutrophils, identified by CD15^+^CD66b^+^citH3^+^ staining (Figure 1A, upper panel and Figure 1B, 2.4 ± 0.97 % and 1.6 ± 1.22 % positive cells, respectively, Figure S1A). Addition of ionomycin strongly induced NET formation, as demonstrated by a significant increase of citH3-positive cells (54.52 ±14.41%) after two hours of incubation (Figure 1A, bottom panel and Figure 1B). This effect was almost completely abolished by pre-incubation with PBMCsec before ionomycin treatment (3.9 ±1.28% citH3-positive neutrophils). By contrast, vehicle treatment showed only weak reduction of NET-formation (27.94 ±17.71% citH3-positive neutrophils) (Figure 1A, bottom panel, and Figure1 B). As DNA is one of the major constituents of NETs (4), we analysed the amount of extracellular DNA in ionomycin-stimulated samples using cytotox green staining (Figure 1C). After two hours, a only weak cytotox green signal was detected in untreated, PBMCsec- or vehicle treated samples. Whereas stimulation with ionomycin resulted in a drastic increase of extracellular DNA (Figure 1C), addition of PBMCsec almost completely inhibited the release of DNA after ionomycin stimulation. To exclude that the observed effect was due to a direct inhibitory effect of PBMCsec on ionomycin, we also used phorbol 12-myristate 13-acetate (PMA, 100 nM), another well-described inducer of NET formation. As shown in figure 1D, PMA treatment of PBMCsec pre-incubated neutrophils led to a comparable inhibition of DNA release. Additionally, we evaluated the metabolic activity of ionomycin-activated neutrophils (52). Ionomycin-activation resulted in a prominent decrease of metabolic activity which in contrast to vehicle treatment, was completely abolished by the addition of PBMCsec (Figure S1B). Immunostaining of neutrophils for citH3 showed classical NET-structures after ionomycin treatment, which were completely absent in the presence of PBMCsec (Figure 1E). Taken together, these findings indicate that treatment of experimentally activated neutrophils with PBMCsec significantly reduces the formation of NETs.

**Figure 1.**
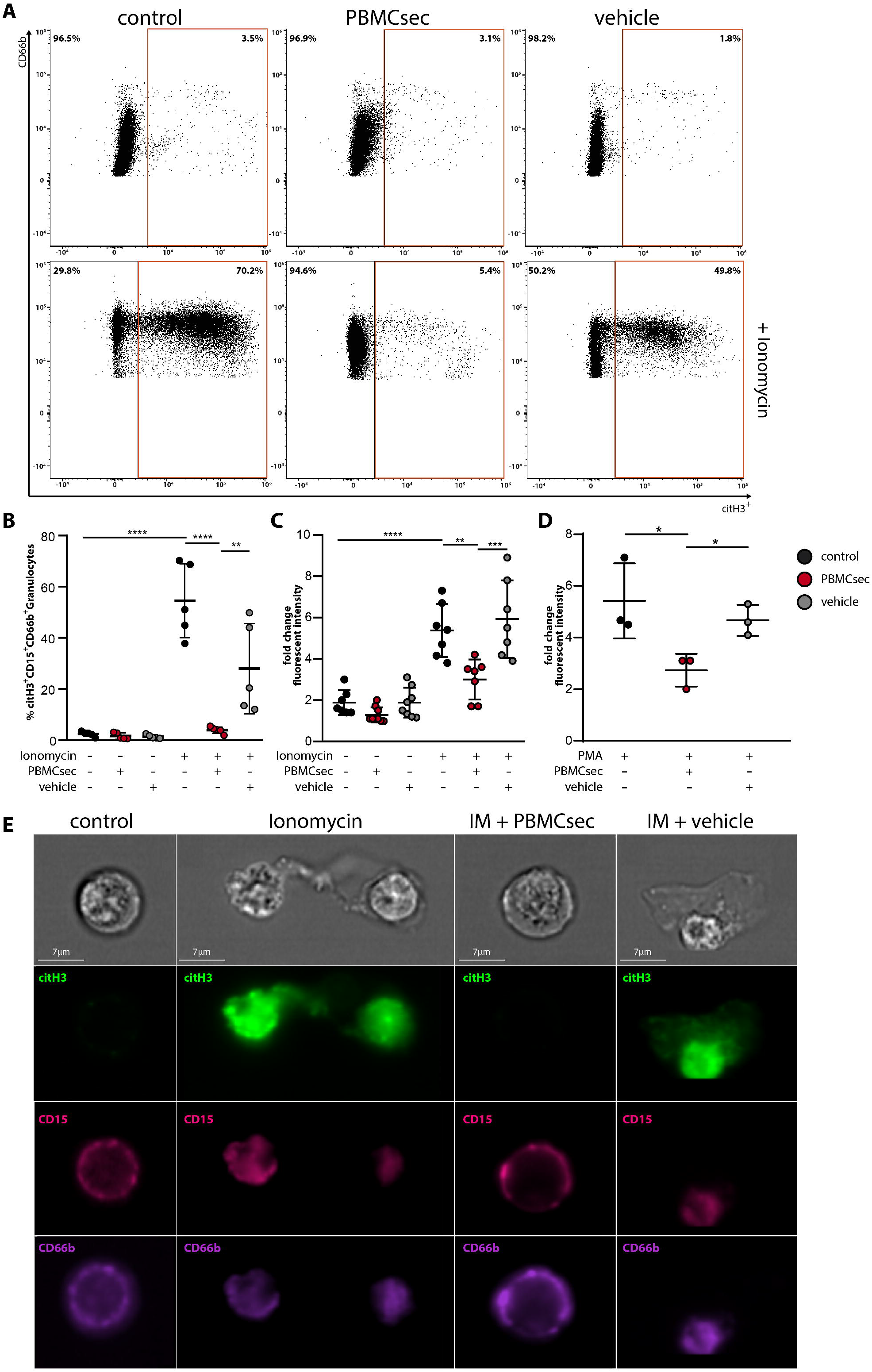
PBMCsec inhibits NET formation. Erythrocyte-lysed blood was treated with PBMCsec or vehicle for 20 minutes and subsequently stimulated with ionomycin (IM) for 2 hours and analysed with flow cytometry, cytotox staining and image stream analysis. **(A)** Neutrophils were identified in flow cytometry as CD66b^+^CD15^+^ cells and NET-forming neutrophils were characterized by citH3. Control, PBMCsec- or vehicle treated samples are shown in the top panel and ionomycin-activated neutrophils are shown in the bottom panel. n = 5, One representative sample is shown out of five replicates summarized in **(B)**. **(C)** Extracellular DNA content was measured using cytotox staining of neutrophils after pre-treatment with PBMCsec or vehicle and subsequent activation for 2 hours with **(C)** IM or **(D)** PMA. Fold change increase of relative fluorescent intensity is shown after two hours of stimulation relative to time point zero/start of stimulation/minute one after induction of NETs. **(E)** Visualization of IM-activated neutrophils was performed using image stream analyses. Untreated (control) and IM-PBMCsec treated neutrophils did not show citH3^+^ staining. IM and IM-vehicle treated neutrophils showed robust citH3^+^ staining of cells and additional extracellular structures (indicative for NETs). Green, citH3; magenta, CD15; purple, CD66b; n = 2, One representative sample is shown. Data are represented as individual values with mean and error bars indicate SD, one-way ANOVA and Sidak’s multiple comparisons test. *p<0.0332, **p<0.0021, ***p<0.0002, ****p<0.0001

### A synergistic effect of different substance classes inhibits NET formation

PBMCsec is composed of different substance classes, including free DNA, lipids, proteins and extracellular vesicles (41,42,50,53) (Figure 2A). Thus, we further aimed to investigate to what extent the individual fractions contribute to the inhibitory effects on NET formation (Figure 2B, C). Therefore, PBMCsec and its fractions were added to whole blood prior to ionomycin stimulation and citH3 levels were analysed (Figure 2B and C, Figure S2). Ionomycin treatment showed a significant increase in citH3^+^ neutrophils, which was almost completely abolished by the addition of PBMCsec. In contrast, purified fractions showed only partial inhibition of ionomycin-induced histone citrullination, indicating that the inhibitory effect of PBMCsec requires the complex interplay of all fractions of the secretome.

**Figure 2.**
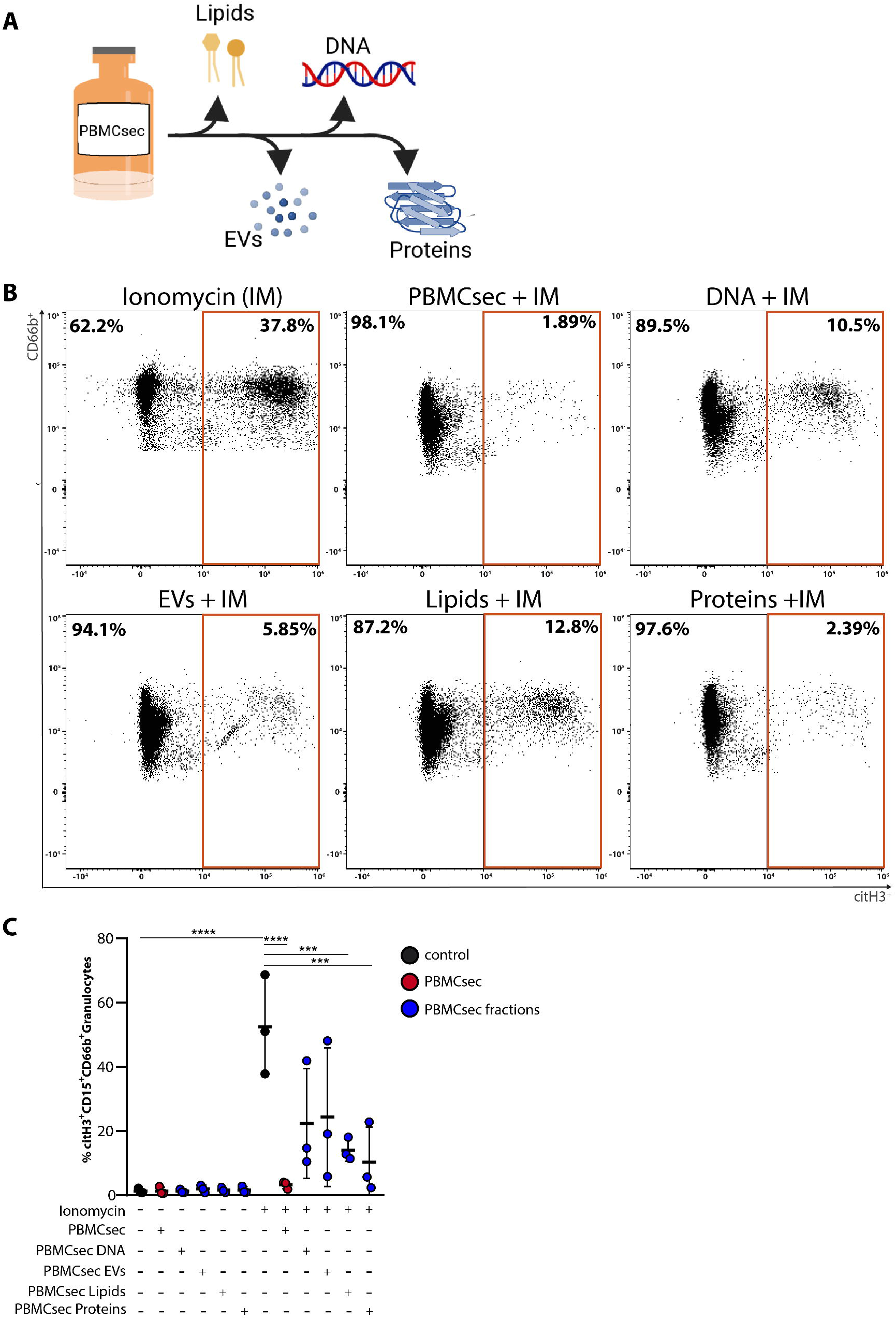
Isolated substance classes of PBMCsec show a synergistic effect on NET-inhibition. **(A)** Schematic depiction of the isolated and tested substance classes of PBMCsec. This scheme was created with BioRender.com**(B)** Neutrophils were identified in flow cytometry as CD66b^+^CD15^+^ cells and NET formation was characterized by citH3^+^ signal. n = 3. Data are represented as individual values with mean and error bars indicate SD. One representative sample out of three is shown, summarized in **(C)** One-way ANOVA and Sidak’s multiple comparisons test. *p<0.0332, **p<0.0021, ***p<0.0002, ****p<0.0001

### PBMCsec does not show DNase-activity

As digestion of NETs by DNAses is the main NETs-clearing mechanism (54), we next investigated whether PBMCsec displays DNAse activity. Therefore, we incubated λDNA with PBMCsec and analysed DNA degradation (Fig 3A). Compared to recombinant DNAse I, which completely digested λDNA, PBMCsec showed no DNA degrading activity (Figure 3A). Since this *in vitro* assay was optimized for DNAse I only, we further tested potential NETs-degrading properties of PBMCsec in whole blood *ex vivo*. For this purpose, we stimulated whole blood with ionomycin and applied PBMCsec either prior to or two hours after ionomycin treatment (Fig 3B). In contrast to neutrophils treated with PBMCsec prior to their activation, treatment two hours after induction of NET formation did not reduce the amount of citH3 positive neutrophils (Figure 3C, D, Figure S3). These data demonstrate that PBMCsec does not degrade pre-formed NETs by DNases, suggesting an active intervention in the NET-forming signalling cascade.

**Figure 3.**
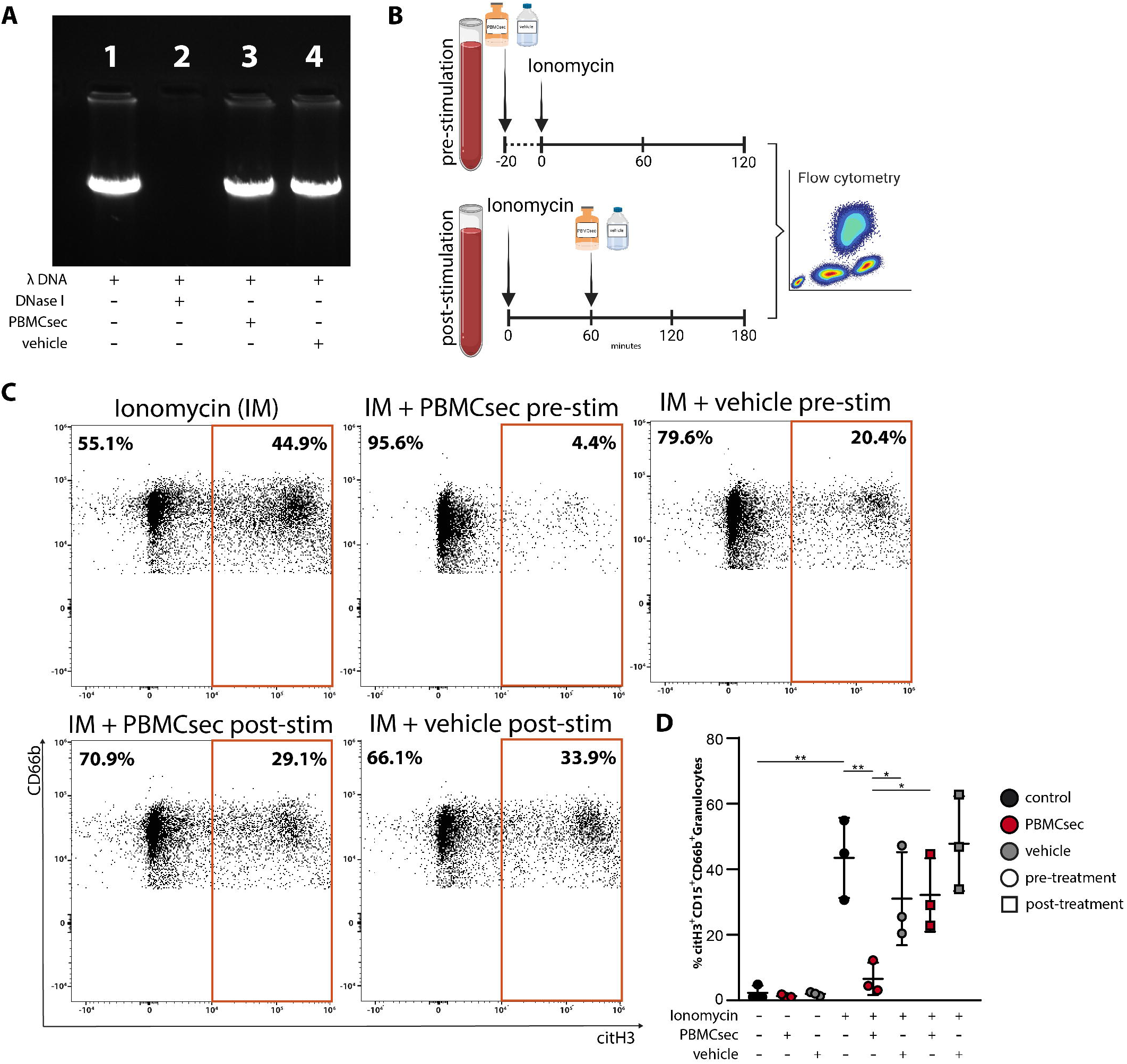
PBMCsec inhibits NET formation by a DNase-independent mode of action. **(A)** DNase activity was measured in a cell-free assay by co-incubation of PBMCsec or vehicle with λ-DNA. DNase I was used as positive control. n = 3, one representative sample is shown. **(B)** Schematic depiction of the adapted neutrophil stimulation protocol for the measurement of potential DNase activity in a cell-based assay. This scheme was created with BioRender.com **(C)** Neutrophils were identified as CD66b^+^CD15^+^ cells and citH3^+^ signal was used to characterize NET formation. n = 3, one representative experiment is shown. **(D)** Statistical summary of all biological donors is shown. Data are represented as individual values with mean and error bars indicate SD. One-way ANOVA and Sidak’s multiple comparisons test. *p<0.0332, **p<0.0021

### PBMCsec inhibits NET formation by preventing ROS production and PAD4 activity

Induction of NETs requires an increase in intracellular calcium levels (12,13,15). We therefore first investigated whether PBMCsec interferes with ionomycin-induced calcium influx. Analysis of intracellular calcium signalling, using a Fura Red based flow cytometry approach, revealed that pre-treatment of whole blood with PBMCsec only marginally reduced calcium influx after addition of ionomycin (Figure 4A and B). The decline in calcium flux was only transient and returned to control values rapidly. In addition, no significant difference was observed between PBMCsec and vehicle treatment, suggesting that the observed decrease of the calcium influx is not sufficient to affect NET formation. As increased intracellular calcium concentrations promote the activity of PKC (13), we next investigated PKC phosphorylation, and were not able to detect differences in the amount of phosphorylated PKC after pre-treating neutrophils with PBMCsec or vehicle (data not shown).

**Figure 4.**
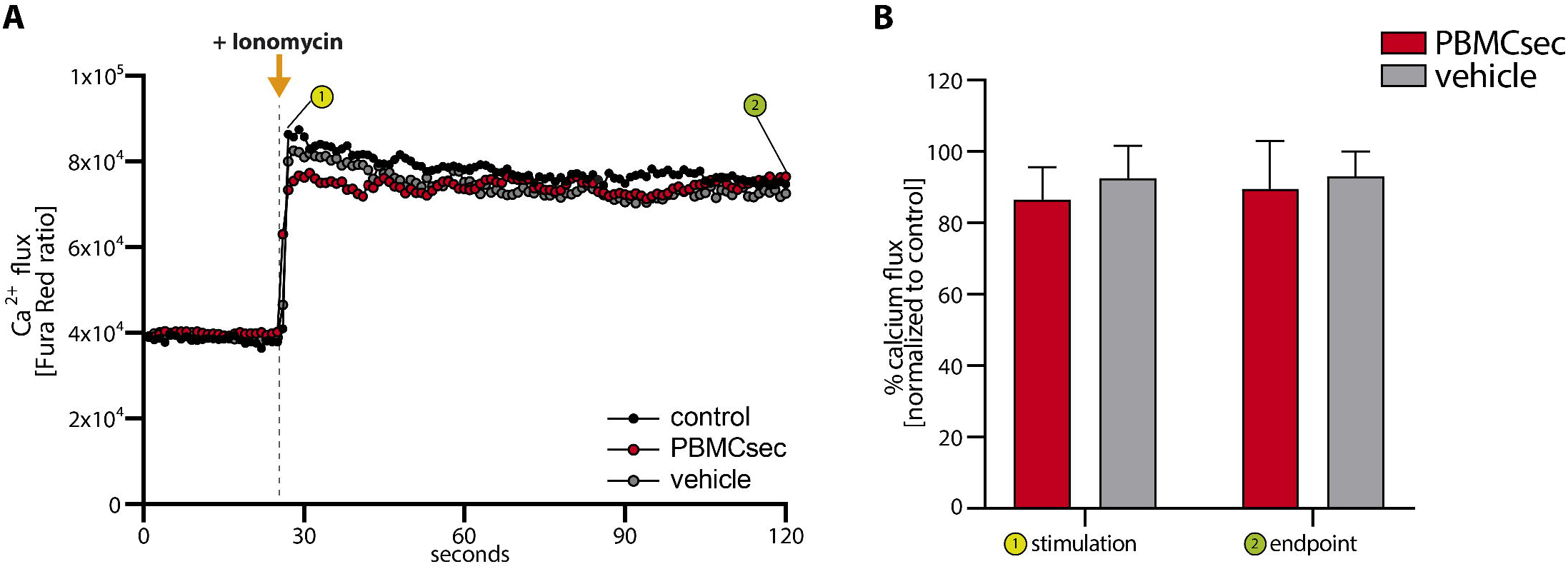
PBMCsec inhibits NET formation without interfering with calcium flux. **(A)** Ratiometric calcium flux was measured with Fura Red in neutrophils. Neutrophils were observed for approximately 30 seconds to record a baseline Fura Red ratio indicating homeostatic calcium flux prior to the addition of IM and subsequent analysis for a total of 120 seconds. **(B)** Statistical analysis of the percent reduction of calcium flux in PBMCsec or vehicle treated neutrophils compared to control samples is shown for the time points of stimulation (①, addition of IM) and the endpoint (②,120 seconds). Error bars indicate SD. No statistical significant reduction was observed for PBMCsec or vehicle treatment.

Since activation of the NET signalling pathway down-stream of NADPH leads to the production of ROS and activation of PAD4 (12,14,15), we next investigated whether PBMCsec exerts its inhibitory activity by modulating these processes. We therefore investigated ionomycin-induced production of ROS using the cell permeant reagent 2’,7’ –dichlorofluorescin diacetate (DCFDA). Ionomycin treatment of PBMCsec-stimulated neutrophils resulted in a significant decrease in ROS production, as compared to ionomycin treatment alone (Figure 5A). By contrast, pre-treatment with vehicle showed no inhibitory effect (Fig 5A). Analysis of protein expression revealed that PBMCsec inhibited ionomycin-induced down-regulation of known anti-oxidative factors, including hemoxygenase-1 (HO-1) (55) and hypoxia inducible factor 1 alpha (HIF-1α) (56) in purified human neutrophils (Figure 5B), which was not observed in vehicle-treated cells.

**Figure 5.**
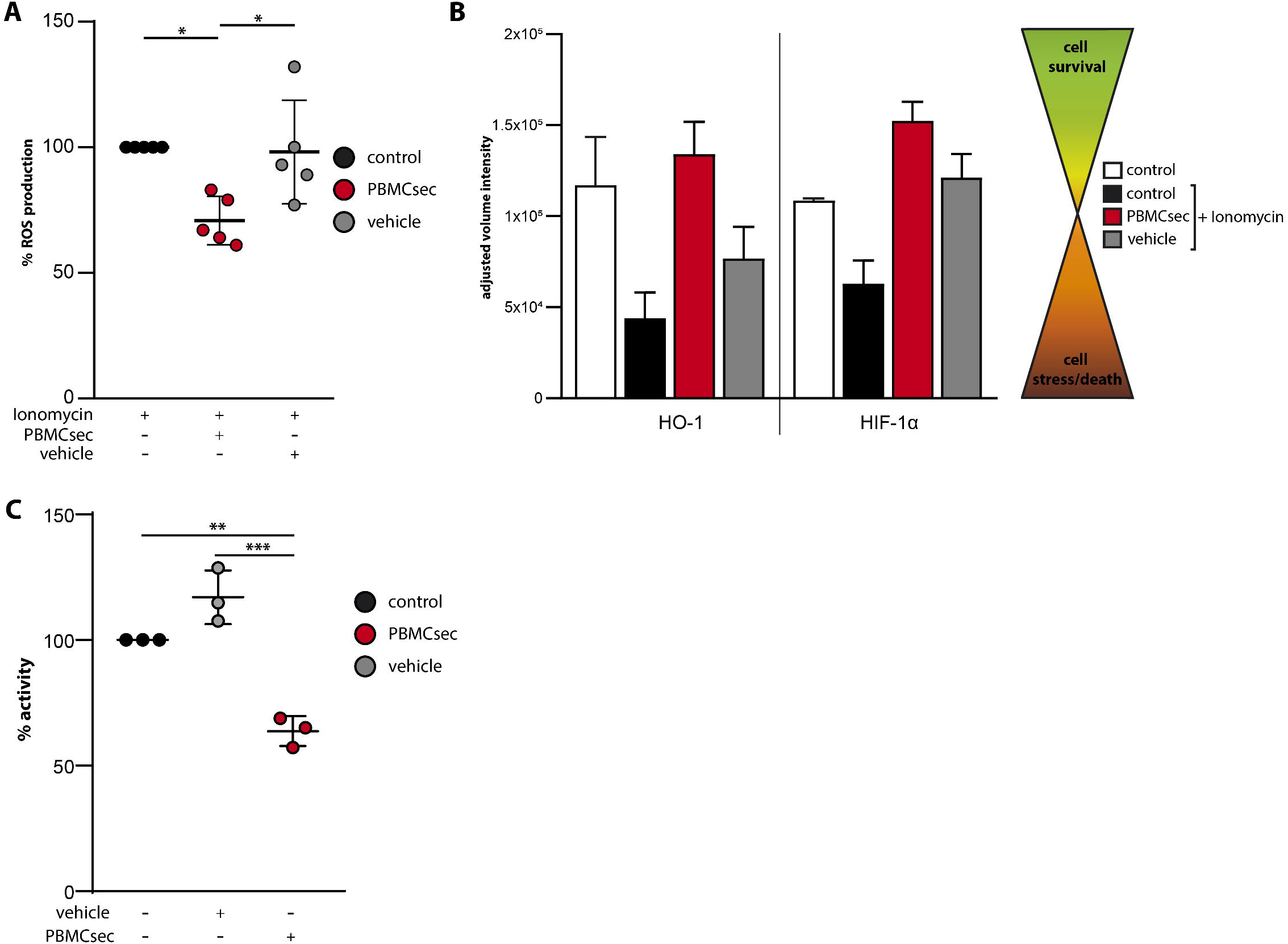
PBMCsec prevents ROS production and PAD4 activity. **(A)** ROS production in IM stimulated and PBMCsec or vehicle treated neutrophils normalized to IM-induced ROS production. n = 5 **(B)** Analysis of protein levels of HO-1 and HIF-1α of isolated neutrophils stimulated with ionomycin and treated with PBMCsec or vehicle, using a proteome profiler is shown. Cell lysates of four individual donors and experiments were pooled. Error bars indicate SD of two technical replicates. **(C)** Enzymatic activity of PAD4 was measured in a cell-free assay upon co-incubation of PBMCsec or vehicle with the PAD4-substrate and normalized to untreated control. n = 3, Data are represented as individual values with mean and error bars indicate SD, unless indicated otherwise. One-way ANOVA and Sidak’s multiple comparisons test was performed where significances are indicated. *p<0.0332, **p<0.0021, ***p<0.0002

The activation of PAD4 with subsequent histone-citrullination represents one of the final steps in the NETs signalling pathway (15). Thus, we next investigated PAD4 activity in a cell-free assay. While vehicle treatment did not show PAD4 inhibiting properties, PBMCsec inhibited PAD4 activity by approximately 40% (Figure 5C). Together, these data demonstrate that PBMCsec exerts its inhibitory activity on NET formation by reducing intracellular ROS production and preventing PAD4 activation.

## Discussion

The formation of NETs is a highly effective first line defence mechanism against invading pathogens (21). However, there is growing evidence that excessive NETs formation contributes to tissue damage and the induction of auto-immune diseases (20,57–59). Although several substances, including acetylsalicylic acid, cyclosporine A (60), metformin (61) and chloroquine (62), have been shown to influence NET formation, therapeutic drugs targeting NET formation are so far not available. In the current study, we provide evidence that PBMCsec effectively inhibited NET formation by reducing ROS production and PAD4 activation, thereby providing a novel potential cell-derived but cell-free therapeutic intervention for NET-associated diseases.

The tissue regenerative and anti-inflammatory action spectrum of PBMCsec is multifaceted (33,35,38,41,47,49,50), and most of its beneficial effects have been shown to require the interplay of several components of the secretome (34,63). Indeed, we also found that NET formation was only fully inhibited when neutrophils were treated with the whole secretome. Since we observed NETs inhibition at different steps of the NETs signalling pathway, we hypothesize that individual secretome fractions act on different signalling molecules. Recently, Laggner et al. showed that lipids present in PBMCsec attenuate skin inflammation and allergic reactions by targeting dendritic cell function (41) as well as mast cell and basophil activation, respectively (submitted manuscript), suggesting that lipids are mainly responsible for the anti-inflammatory activities of PBMCsec. Several lipid species have been detected in PBMCsec, including phosphatidylserines, lysophosphatidylcholines, lysophosphatidylethanolamines, phosphatidylcholines, phosphatdiylethanolamines and resolvins (41). Interestingly, several studies described a NET-inhibitory or – resolving action of resolvins (64–66). Spinosa et al. demonstrated a decreased NET burden accompanied by reduced abdominal aortic aneurysm in reslovin-treated mice (64). In addition, neutrophils derived from resolvin-treated mice showed less susceptibility to ionomycin-induced NET formation (65). While both of these studies identified less NET formation in the presence of resolvins, Chiang et al. showed that NETs formed after *Staphylococcus aureus* infection were more efficiently cleared by macrophages after treatment with reslovins (66). These data indicate an important function of resolvins in the prevention of NET formation and/or resolution of NETs. Therefore, it is conceivable that the partial inhibition of NET formation observed by PBMCsec-lipids may be explained by the variety of resolvins found in PBMCsec (41). However, whether PBMCsec-derived resolvins or other lipid classes are indeed involved in PBMCsec-induced NETs inhibition needs to be determined in future studies.

In addition to the lipid fraction, PBMCsec-derived proteins also showed a strong inhibitory action on NET formation. Previous studies demonstrated that addition of either bovine or human serum albumin to ionomycin-treated cells almost completely blocked NET formation by chelating calcium (67). However, as we only detected a slight decrease in calcium influx after treatment with PBMCsec and vehicle, a sole albumin-dependent effect is unlikely. Furthermore, we also showed comparable effects when NET formation was induced with PMA, which induces NETs in a calcium-independent manner. Together, our data suggest a calcium- and albumin-independent mode of action of the protein fraction of PBMCsec. Further in-depth proteomics analyses of PBMCsec-derived proteins are required to elucidate whether a single protein or a combination of proteins is responsible for the inhibition of NET formation.

Our data suggest the inhibition of ROS production and PAD4 activation as the two major modes of action for the reduction of NET formation by PBMCsec. Oxidative stress, especially the generation of ROS, is a hallmark of NET formation (14,68) and HSPs are known to effectively block excessive ROS production (69). Our study revealed that PBMCsec inhibited hemeoxygenase 1 (HO-1 or HSP32) and HIF-1α downregulation during ionomycin-induced NET formation. Both, HO-1 and HIF-1α, have been shown to promote neutrophil survival by reducing ROS levels (70) and via Akt and NFκB signaling under stress, respectively (56,71). These data suggest that PBMCsec contributes to the stabilization of the delicate balance of pro- and anti-oxidative processes by regulating the expression of HSPs, thereby preventing neutrophil-induced tissue damage. Since HO-1 is also known to downregulate adhesion molecules and chemokines required for neutrophil infiltration (55), PBMCsec may alleviate inflammatory responses by reducing neutrophil infiltration in damaged and inflamed tissue.

PAD4 is one of the most prominently investigated factors critical for NET formation, and PAD inhibitors have been extensively studied in the context of a broad variety of diseases, including multiple sclerosis (72), myocardial infarction (73) and rheumatoid arthritis (74). However, the exact mechanism of PAD4 inhibition is not yet fully understood (74). Our data indicate that stressed PBMCs secrete factors that serve as PAD4 inhibitors. Interestingly, Yost et al. identified a group of peptides in umbilical cord blood with strong PAD4-inhibiting effects, leading to inhibition of NETs (75). Sequence analyses identified α1-antitrypsin, a serine protease inhibitor, known to possess immunomodulatory and anti-inflammatory properties (76), as the main PAD4-inhibiting factor. Interestingly, α1-antitrypsin is synthesized by circulating monocytes and therefore a component of PBMCsec (Figure S5) (77). This enzyme inhibitor has been considered as acute phase protein, which contributes to inhibition of NET formation by targeting a vast array of factors NET formation (78). Further studies are needed to identify the PAD4-inhibiting factor(s) in PBMCsec.

In summary, we have demonstrated a strong NETs-inhibitory activity of PBMCsec via a dual mechanism. Specifically the identification of a PAD4 inhibitor, produced naturally in the human body, might strongly improve the treatment of diseases associated with excessive NET formation, such as rheumatoid arthritis (74), multiple sclerosis (72), sepsis (21), heart failure and myocardial infarction (73). Pre-clinical toxicological evaluation of PBMCsec has already been performed without the occurrence of major adverse events after topical and intravenous application (88, LPT, study number 35015). Therefore, our study has paved the way for a clinical study in humans, assessing the potency of PBMCsec in NETs-associated diseases *in vivo*.

## Materials and Methods

### Ethics statement

This study was conducted in accordance with the Declaration of Helsinki and applicable local regulations. Use of human neutrophils was approved by the institutional ethical review board of the Medical University of Vienna (Vienna, Austria). Written informed consent was obtained from all donors.

### Generation of PBMCsec

PBMCsec was produced in compliance with good manufacturing practice by the Austrian Red Cross, Blood Transfusion Service for Upper Austria (Linz, Austria) as previously described (34,41). Briefly, PBMCs were enriched using Ficoll-Paque PLUS (GE Healthcare, Chicago IL, USA) density gradient centrifugation. Cell suspensions were adjusted to 2.5 x 10^7^ cells / mL and exposed to 60 Gy γ-irradiation (IBL 437, Isotopen Diagnostik CIS GmbH, Dreieich, Germany). Subsequently, cells were cultured in phenol red-free CellGenix GMP DC medium (CellGenix, Freiburg, Germany) for 24 hours. Cells as well as cellular debris were removed by centrifugation. The conditioned supernatants containing the secretome were filtered through 0.22 μm filters followed by viral clearance using Theraflex methylene blue technology (MacoPharma, Mouvaux, France). The secretomes were lyophilized and sterilized by high-dose γ-irradiation (25 000 Gy, Gammatron 1500, Mediscan, Seibersdorf, Austria). CellGenix GMP DC medium without cells was used as vehicle control. The GMP batches A000918399131, A00918399136 and A000918399132 were used in this study. The stock concentration of one vial lyophilized secretome equals to 25 units/mL.

### Fractionating PBMCsec

The lipid fraction was purified as previously described by Laggner et al. (41). The protein fraction was isolated by combining four times the volume of ice cold acetone (VWR Chemicals, PA, USA) to one volume of reconstituted PBMCsec, followed by thorough vortexing and incubation at −20 °C for 60 minutes. To obtain a protein precipitate, the sample was centrifuged at 18 000 g for 10 minutes. To obtain the protein-free PBMCsec, the liquid phase was lyophilized and subsequently resuspended in 0.9 % NaCl at the same volume as initially used and stored at −20 °C until further use. To further purify the protein fraction, ice-cold acetone was added to the protein pellet, briefly vortexed and centrifuged at 18 000g for 10 minutes. Acetone was discarded and remaining acetone was allowed to evaporate at room temperature. Finally, the protein pellet was resuspended in 0.9 % NaCl in the initially used volume. DNA was isolated by adding equal amounts of isopropanol (Merck Millipore, MA, USA) as PBMCsec and 1/10 volume of 7.5 M sodium acetate (Merck Millipore) and incubated at −20°C for 1 hour. After centrifugation for 5 minutes at 18 000 g the DNA pellet was washed twice with one mL 70% Ethanol (Merck Millipore) followed by 5 minutes centrifugation at 18 000g. The DNA pellet was allowed to dry for 10 minutes at room temperature prior to resuspension in double distilled, nuclease-free H_2_O. Extracellular vesicles were obtained by ultracentrifugation at 110 000 g for 2 hours at 4 °C as previously described (34). To ensure comparability, all fractions were used in the same concentrations as present in PBMCsec.

### Neutrophil isolation

Neutrophils were isolated using the MACSxpress Whole Blood Neutrophil Isolation kit (Miltenyi Biotec, Bergisch-Gladbach, Germany) according to manufacturer’s instructions. In brief, magnetic beads were resuspended in 2 mL of buffer A. One fourth of the total amount of processed blood of magnetic beads and buffer B were added to the blood and incubated at room temperature for 5 minutes under constant gentle rotation. Blood and isolation cocktail mix were placed in the MACSxpress Separator (Miltenyi Biotec) and allowed to separate for 15 minutes. Clear, neutrophil-containing top phase was transferred into a fresh tube and washed with basal RPMI 1640 without phenol red (Thermo Fisher Scientific, Waltham, USA). If required, a red blood cell lysis was performed using Red Blood Cell Lysis Buffer (Abcam, Cambridge, UK) for 10 to15 minutes at room temperature. Neutrophils were resuspended in basal RPMI 1640 in an assay dependent concentration without phenol red for further use.

### Induction of NET formation

Either isolated neutrophils or whole blood samples after red blood cell lysis were pre-treated with 2 units/mL PBMCsec or equivalent vehicle medium for 20 minutes at 37 °C. Cells were then stimulated with 5 μM ionomycin (Sigma Aldrich, St. Louis, Mo, USA) or 100 nm PMA (Sigma Aldrich) for 2 hours at 37°C unless indicated otherwise.

### Flow cytometry

After stimulation with indicated compounds, cells were centrifuged and stained with anti-citrullinated Histone H3 antibody (ab5103, Abcam) to detect NETs, anti-CD66b antibody (pacific-blue conjugated mouse anti-human, clone G10F5, BioLegend, San Diego, CA, USA) and anti-CD15 antibody (phycoerithrin-cyanine 7 conjugated mouse anti-human, clone W6D3, BioLegend) to identify neutrophils. Flow cytometric analysis was performed using BD FACSCanto II and BD FACSDiva software (version 6.1.3) (BD Pharmingen, San Jose, CA, USA).

### Cell viability assay

Incucyte Cytotox Dye for Counting Dead Cells (Sartorius, Goettingen, Germany) was used according to manufacturer’s instructions. In brief, cells were treated as indicated followed by the addition of 250 nM cytotox green dye for staining 100 μL cell suspension in a concentration of 4×10^6^ cells/mL condition in a 96-well plate. Cell death was assessed over the indicated time periods in a microplate reader at an excitation wavelength of 485 nm and an emission wavelength of 520 nm. Microplate reader (BMG Labtech, FLUOstar OPTIMA) and the BMG Labtech Optima software (software version 2.20Rs).

### EZ4U cell proliferation and cytotoxicity assay

EZ4U (Biomedica, Vienna, Austria) assay was performed according to manufacturer’s instructions. Briefly, the substrate was dissolved in 2.5 ml activator solution and pre-warmed to 37 °C. 20 μL substrate was added to 200 μL cell suspension at a concentration of 4×10^6^ cells per condition in a 96-well plate and incubated for 2 hours. Continuous absorbance measurements at 450 nm were performed using a microplate reader (BMG Labtech, FLUOstar OPTIMA) and the BMG Labtech Optima software (software version 2.20Rs).

### ROS production measurement

ROS production was measured using the DCFD/H2DCFDA cellular ROS assay kit (Abcam). Cells were treated as indicated and the assay was performed as recommended by the manufacturer.

### Ca^2+^ flux measurement

Ratiometric calcium flux measurements with Fura Red were performed as described by Wendt et al. with minor modifications (80). In Brief, a cell suspension of 4 x 10^6^ cells per condition, either isolated neutrophils or whole blood, pre-treated with PBMCsec or vehicle for 20 minutes as indicated, were washed, resuspended in 400 μL full medium containing 1 μM Fura Red (Invitrogen, Thermo Fisher Scientific, Waltham, MA) and incubated for 30 minutes at 37 °C. Cells were washed once with medium, resuspended in 4 mL medium and incubated for another 30 minutes at 37 °C. Subsequently, cells were rested on ice for up to 30 minutes. Data was acquired on a FACSAria III flow cytometer (BD Bioscience). Before intracellular calcium flux measurement, 1 mL of Fura Red-loaded cells was transferred to a FACS tube and pre-warmed for 5 minutes at 37 °C in a water bath. The cells were kept at 37 °C during the whole measurement. The baseline response was recorded for 30 seconds prior to stimulation with 5 μM ionomycin. Changes in calcium mobilization were recorded for a total of 120 seconds. Fura Red was excited using a 405 nm violet laser and a 561 nm green laser and changes in emission were detected with a 635LP, 660/20 BP and a 655LP, 795/40 BP filter set, respectively. The ‘Fura Red Ratio’ over time was calculated using the Kinetics tool in FlowJo software (version 9.3.3, Tree Star Inc., Ashland, OR, USA) as follows:

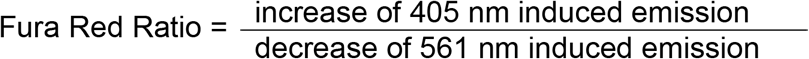

### DNase activity measurement

DNase activity was measured by incubating 0.25 μg/μL Lambda DNA with 0.5 M acetate/NaOH at pH 4.8 (Merck Millipore), 50 mM CaCl_2_(Merck Millipore), 50 mM MgCl_2_ (Sigma-Aldrich, St. Louis, USA), 40 mM 2-mercaptoethanol (Merck Millipore) and either 2 units/mL PBMCsec or equivalent vehicle control or DNase I (Thermo Fisher Scientific) as positive control for 1 hour at 37°C. 10 μl of each sample were loaded into a 1% agarose gel with gel red (Biotium, Fremont, CA, USA) together with 2 μL 6x loading dye (Thermo Fisher Scientific). Electrophoresis was performed at 200 V for 35 minutes. DNase activity was determined by absence of λ-DNA.

### Proteome Profiler

Human Apoptosis Array kit (R&D Systems, Minneapolis, MN, USA) was performed in accordance to manufacturer’s instructions with no modifications. Isolated neutrophils were treated as indicated and cell lysates of 4×10^6^ cells per condition of 4 individual donors were pooled.

### PAD4 inhibitor assay

Inhibitory capacity of PBMCsec was measured using the PAD4 inhibitor Screening Assay kit (ammonia, Cayman Chemical, Ann Arbor, MI, USA) and performed according to manufacturer’s instructions.

### Statistics

Statistical analyses were performed using Prism 8.0.1 (Graph Pad Prism). Data was shown as ± standard deviation (SD). One-way ANOVA and Sidak’s multiple comparisons test were performed and *p<0.0332, **p<0.0021, ***p<0.0002, ****p<0.0001.

## Supporting information

Supplementary Figures and graphical abstract

## Abbreviations

DCFDA: 2’,7’-dichlorofluorescin diacetate
HIF-1α: hypoxia inducible factor 1 alpha
HO-1: hemeoxygenase-1
HSP: heat shock protein
IM: ionomycin
NET: neutrophil extracellular trap
PAD4: peptidyl arginine deiminase 4
PBMC: peripheral blood mononuclear cells
PBMCsec: secretome of stressed peripheral blood mononuclear cells
PKC: protein kinase C
PMA: phorbol 12-myristate 13-acetate
ROS: reactive oxygen species

## Author Contributions

Conceptualization, KK, MM and HJA; methodology, KK, MM., ASO and TMH; validation KK, MM, ASO, TMH; formal analysis, KK; investigation, KK, ASO, TMH, KP; resources, HJA.; data curation, KK, DC, MD, DB; writing—original draft preparation, KK, MM; writing—review and editing, KK, MM, ML, AM; visualization, KK; supervision, MM; project administration, KK, MM; funding acquisition, HJA. All authors have read and agreed to the published version of the manuscript.

## Acknowledgements

We thank HP Haselsteiner and the CRISCAR Familienstiftung for their belief in this private–public partnership to augment basic and translational clinical research.

