## Supplementary Figures and graphical abstract for "The effect of paracrine factors released by irradiated peripheral blood mononuclear cells on neutrophil extracellular trap formation": 2022-05-23_Supplementary figures + graphical abstract_Paracrine factors released by....pdf

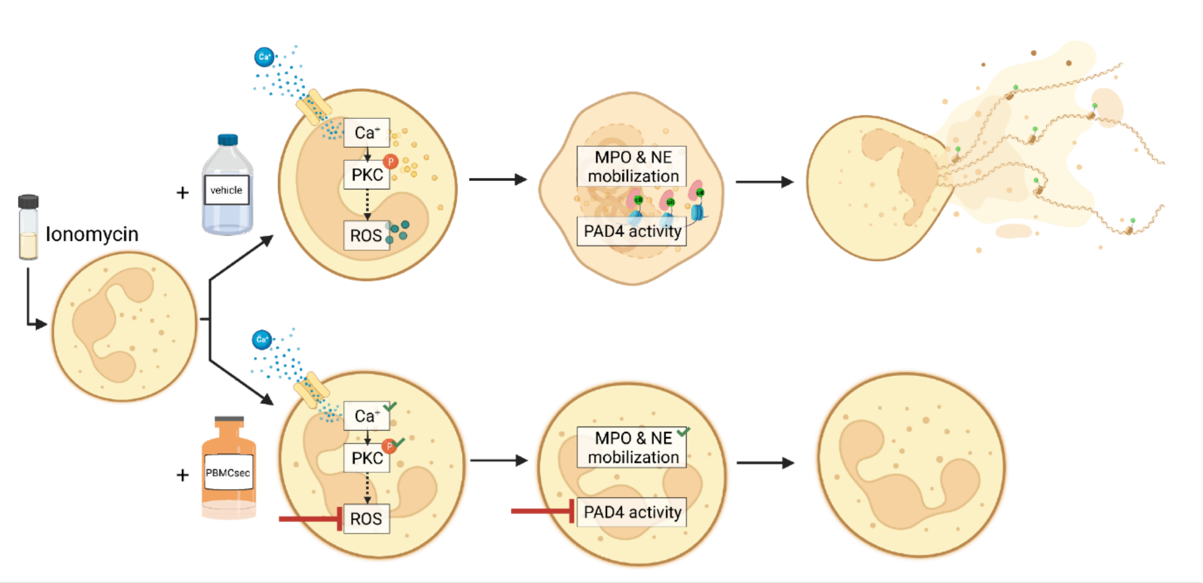


*Graphical overview*

Ionomycin (IM) treatment of neutrophils initiates a series of intracellular events driving neutrophils into NETosis. Intracellular calcium concentrations increase by tremendous influx through membrane channels and liberation of intracellular calcium from the endoplasmic reticulum storage. Protein kinase C (PKC) is phosphorylated and induces several downstream events, depending on the form of NET formation, leading to increased reactive oxygen species (ROS) production which facilitates myeloperoxidase (MPO) and neutrophil elastase (NE) mobilization as well as enzymatic activity. This further promotes peptidyl arginase deiminase 4 (PAD4) activity resulting in histone citrullination and eventually the extrusion of NETs. Vehicle treatment (upper panel) does not interfere with IM-induced NET formation. PBMCsec treatment of IM-activated neutrophils blocks ROS production and PAD4 activity thereby tremendously impairing major components of the NET-signalling cascade resulting in inhibition of NETosis. Created with BioRender.com

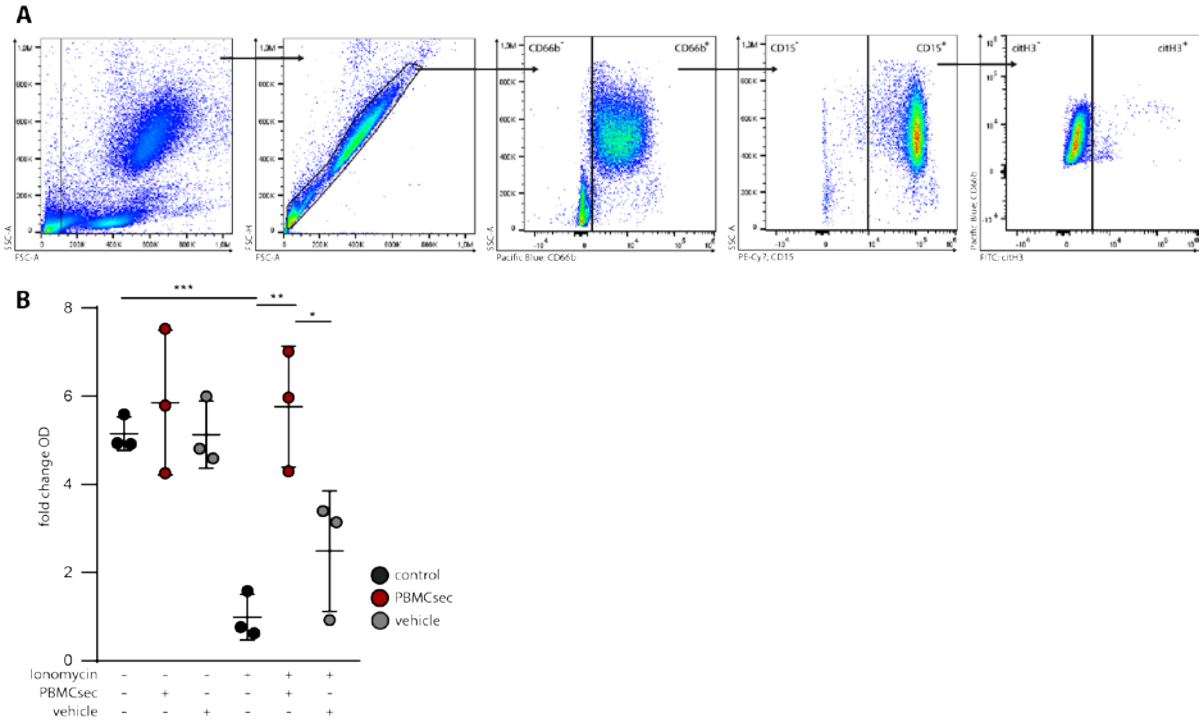


*Figure S1*

(A) Flow cytometry gating strategy for erythrocyte lysed blood samples is shown. (B) Metabolic activity of neutrophils was measured using an absorbance based assay (EZ4U). Vehicle treated neutrophils did not show altered metabolic activity compared to untreated control samples. PBMCsec treatment appeared to partially promote metabolic activity of non-activated neutrophils. IM treatment resulted in a significant reduction of metabolic activity of neutrophils which was abolished by PBMCsec treatment. Vehicle treatment could not restore homeostatic metabolic activity in IM-activated neutrophils. Data are represented as mean ± SD, one-way ANOVA and Sidak's multiple comparisons test. *$p<0.0332$, **$p<0.0021$, ***$p<0.0002$

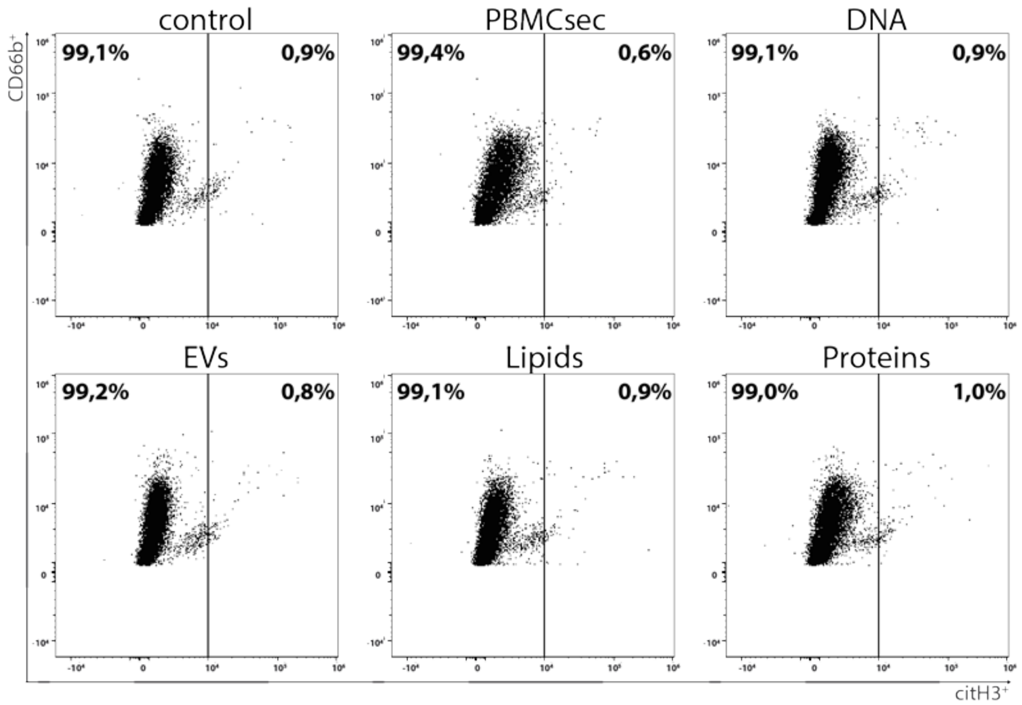


*Figure S2*

Flow cytometric analysis of untreated control neutrophils and neutrophils treated with PBMCsec derived substance classes in absence of an activating stimulus is shown. Neutrophils were identified as $CD66b^{+}CD15^{+}$ cells and NET formation was characterized by additional $citH3^{+}$ signal. n = 3, one representative experiment is shown.

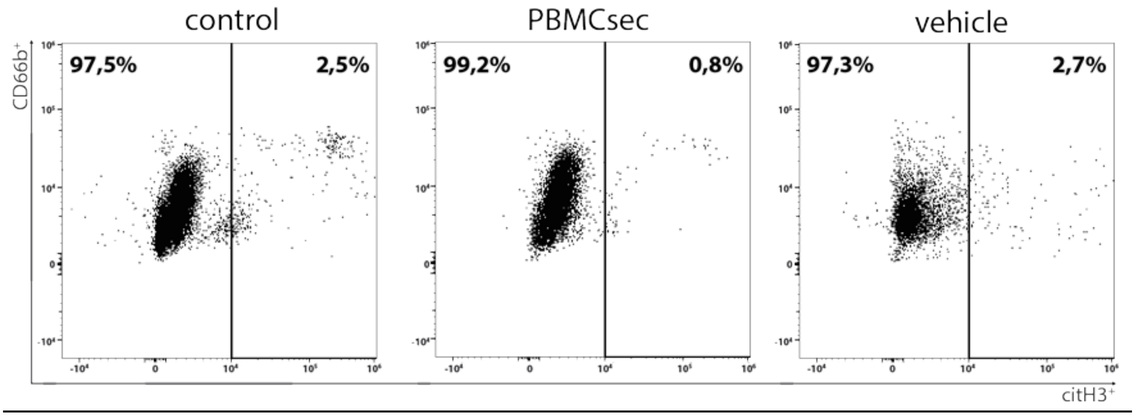


*Figure S3*

Flow cytometric analysis of untreated control neutrophils and neutrophils treated with PBMCsec or vehicle in absence of an activating stimulus after two hours is shown. Neutrophils were identified as $CD66b^{+}CD15^{+}$ cells and NET formation was characterized by additional $citH3^{+}$ signal. n = 3, one representative experiment is shown.

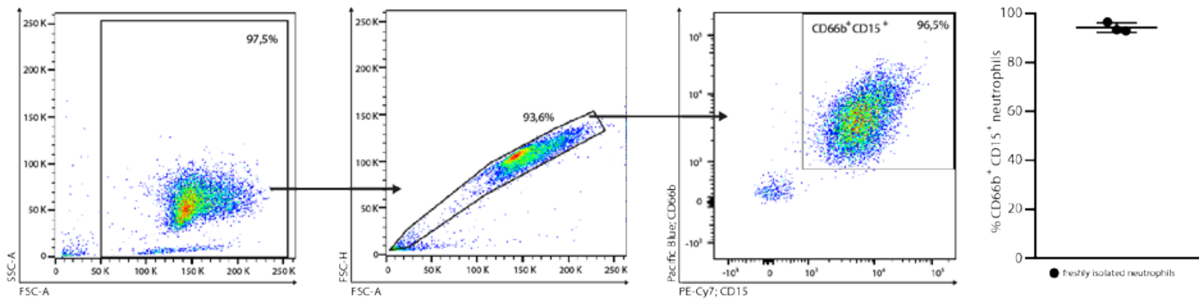


*Figure S4*

Flow cytometry gating strategy for the analysis of neutrophil purity after magnetic bead isolation is shown. Neutrophil purity was assessed by the percentage of CD66b⁺CD15⁺ cells and ranged from 92.9% to 96.5%.

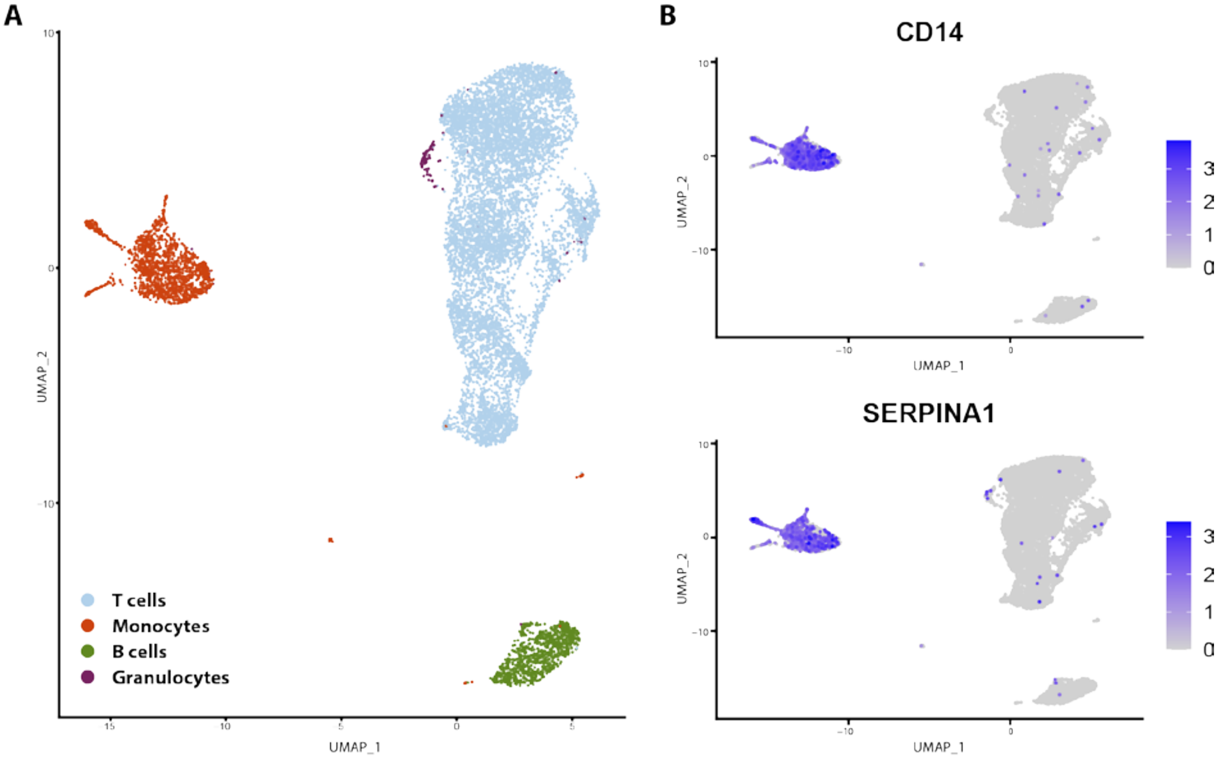


*Figure S5*

Single cell RNA sequencing analysis of erythrocyte lysed whole blood shows (A) UMAP cluster depiction of captured cell populations. (B) Monocyte cluster were identified by the expression of CD14. (C) SERPINA1 expression was almost exclusively found in the $CD14^{+}$ monocyte cluster.
